# A Bimolecular Multicelular complementation system for the detection of syncytium formation: A new methodology for the identification of entry inhibitors

**DOI:** 10.1101/522540

**Authors:** María Jesús García-Murria, Neus Expósito-Domínguez, Ismael Mingarro, Luis Martinez-Gil

## Abstract

Fusion of viral and cellular membranes is a key step during the viral life cycle. Enveloped viruses trigger this process by means of specialized viral proteins expressed on their surface, the so called viral fusion proteins. There are multiple assays to analyze the viral entry including those that focus on the cell-cell fusion induced by some viral proteins. These methods often rely on the identification of multinucleated cells (syncytium) as a result of cell membrane fusions. In this manuscript, we describe a novel methodology for the study of cell-cell fusion. Our approach, named *Bi*molecular *Mu*lticelular *C*omplementation (BiMuC), provides an adjustable platform to investigate qualitatively and quantitatively the formation of a syncytium. Furthermore, we demonstrated that our procedure meets the requirements of a drug discovery approach and performed a proof of concept small molecule high-throughput screening to identify compounds that could block the entry of the emerging Nipah virus.

## 1. Introduction

Regardless of their entry pathway at some point during their life cycle, viruses must cross the cell membrane. For enveloped viruses, this process requires the fusion of the viral and cellular membranes [1]. Enveloped viruses trigger this membrane merge process by means of specialized viral proteins expressed on their surface, the so called viral fusion proteins. By undergoing intense structural rearrangements viral fusion proteins are capable of lowering the kinetic barriers necessary to achieve the coalition of two biological membranes. Currently three classes of fusion proteins have been characterized according to their structure and mechanism of action: Class I (eg. influenza HA), Class II (represented by flavivirus envelope protein E) and Class III (illustrated by rhabdovirus glycoprotein G). For a detailed review on the characteristics of each viral fusion class visit [1,2]. Fusion can be triggered directly by interactions of the fusion or a companion protein at the viral surface with a cellular receptor on the plasma membrane. In this case, expression of the viral fusion protein together with the attachment protein at the host cell membrane during viral replication can lead to syncytia formation (fusion of neighboring cells generating multi-nucleate cells). Alternatively, the interaction between a viral and cellular protein at the cell surface can prompt particle endocytosis. Subsequently, the low endosomal pH or the interaction with a second internal receptor elicits the fusogenic conformational change required for membrane fusion [2].

Although the vast majority of antivirals focus on viral replication, the key role of fusion and attachment proteins during the virus life cycle makes them an attractive target for therapeutic intervention. There are several successful entry inhibitors in the market (including Human immunodeficiency virus (HIV) and Influenza A virus (IAV) antivirals) and many more in research and development stages [3–8]. There are some advantages of targeting an extracellular protein such as the viral fusion and attachment proteins or the cellular receptors expressed on the cell surface necessary for viral entry. This type of extracellular target sites are much easier to reach for the antiviral, resulting in improved pharmaco-kynetics and lower toxicity profiles of the drug of choice. It would also be advisable to have more of these type of drugs simply to increase our palette of viral inhibitors and therefore the potential therapy combinations, a highly successful treatment regimen extensively used to fight severe infections such those produce by HIV or Hepatits C virus [9,10]. Furthermore, a detailed comprehension of the viral-cellular membrane fusion would trigger the design of new mechanism to block viral entry.

There are multiple assays both *in vitro* and *in vivo* for evaluating the viral entry, including cell-virus fusion assays with pseudotyped viral particles, cell-cell fusion assays, and *in vitro* biochemical assays [11]. Among these, the cell-cell fusion assays, based on the formation of a syncytium, offer a safe and virus free alternative suitable for antiviral development even in a high throughput format. Identification of syncytium formation has traditionally been done by microscopy. These methodologies are far from ideal, they are slow, not quantifiable and lack sensitivity. Furthermore, despite huge advance on image analysis [12,13], the implementation of these methods for high throughput screenings (HTS) is not yet optimal.

Many efforts have been made to facilitate the analysis of cell-cell fusions induced by viral proteins, especially in the HIV field [11,14–19]. Herschhorn et al. [20] described a system based on the fusion of two cell lines, an effector line stably expressing a tetracycline-controlled transactivator (tTA) that controls the expression of HIV-1 Env and a target cell line expressing the HIV-1 receptors CD4 and CCR5 and the firefly luciferase under a tTA-responsive promoter. Env-mediated fusion of these two cell lines grants tTA dependent activation of the F-Luc expression. In an earlier manuscript [21], Bradley J. and colleagues reported a similar methodology using two cell lines, one expressing CD4, CCR5, and the β-galactosidase under a LTR-promoter, while the other expresses constitutively HIV proteins gp160 and tat. Fusion of these two cell lines facilitates the transfer of the tat transcription factor and the accumulation of β-galactosidase. Interestingly, the authors probed that this methodology could be used in a HTS compatible format. All these techniques have proven useful for the development of anti-HIV drugs. However, they are not compatible with other viruses and require the generation of cell lines, which despite having some advantages (mainly the standardization of the results) are time consuming processes and hinder the analysis of mutants and variants of the proteins involved in the process.

In this manuscript we describe a novel methodology for the study of cell-cell fusion. Our approach, named *Bi*molecular *Mu*lticelular *C*omplementation (BiMuC), is based on the bimolecular complementation of a reporter [22] and provides an adjustable platform to investigate the entry processes of multiple viruses. Furthermore, we demonstrated that our procedure meets the requirements of a drug discovery approach and performed a proof of concept small molecule HTS to identify compounds that could block the entry of the emerging Nipah virus (NiV).

## 2. Materials and Methods

### 2.1 Plasmids and cell lines

HEK 293T cells were obtained from ATCC (http://www.atcc.org) and were maintained in Dulbecco’s Modified Eagle Medium (DMEM) (Gibco, http://www.lifetechnologies.com) supplemented with 10% fetal bovine serum (FBS, Gibco).

The NiV F and G plasmids were a gift from Dr. M. Shaw laboratory (Ichan School of Medicine at Mount Sinai) while Hendra virus (HeV) F and G were donated by Dr. BH Lee (Ichan School of Medicine at Mount Sinai). The Jun-Nt VFP and Fos-Ct VFP expressing plasmids were obtained from Addgene (#22012 and #22013 respectively). Finally, the Jun-Nt luciferase and Fos-Ct luciferase plasmids were obtained by substituting the VFP sequence on the Jun and Fos BiFC constructs by the appropriated luciferase segment (obtained from the Promega, pRL_CMV plasmid) [23].

### 2.2 Cell-Cell fusion assay

For the cell-cell fusion assay HEK 293T cells were seeded in 6 well plates (2×10^6^ cells/plate) on day 1 (DMEM supplemented with 10% FBS). After 24 hours (day 2) cells were transfected using polyethylenimine (PEI) as a transfection reagent [24] with either Jun-Nt, Fos-Ct, NiV F and G or mock transfected (1µg of DNA/well). Alternatively, for the 2 pool cell approach one set of cells was transfected with Jun-Nt, NiV F and G while the other received just Fos-Ct. On day 3, cells were counted, mixed into the positive (Jun-Nt, Fos-Ct and NiV F and G expressing cells) and negative controls (Jun-Nt, Fos-Ct and mock transfected cells) and seeded into 24 (1×10^5^ cells/well in 500µl of media), 96 (3×10^4^ cells/well in 100µl of media) or 384 well plates (1×10^4^ in in 25µl of media). In those experiments in which only two cell pools are used, the positive control included cells expressing Jun-Nt, NiV F and G and Fos-Ct while the negative control contained only mock transfected cells. Finally, on day 4. fluorescence was measured directly on the plates (96 and 384 well plates) in a VICTOR™ X Multilabel Plate Reader (Perking Elmer). Alternatively, when using the 24 well plates cells were collected, transferred into a 96 black plate prior to fluorescence measurement. Each experiment was performed at least three times in triplicates. The same protocol was used when the fluorescence reporter was substituted by a luminescence readout in this case we utilize 96 well white cell-culture plates and the Firefly Luciferase Assay kit from Sigma following the manufacturer protocol.

### 2.3 Identification of NiV cell-cell fusion inhibitors

For the identification of NiV F and G inhibitors the afforded mention protocol was adjusted to meet the HTS criteria of the Screening facility at the *Centro de Investigación Principe Felipe* (*CIPF*) (Valencia, Spain). To evaluate the robustness of our assay we calculated the Z’-factor [25] and the Signal-to-Noise (S/N) ratio. Z’-factor=1-((3δ_pos_+3δ_neg_)/(µ_pos_-µ_neg_)), where µ_pos_ is the mean signal for the positive control, µ_neg_ is the mean signal for the negative control, δ_pos_ is the standard deviation of the positive control, and δ_neg_ is the standard deviation for the negative control. S/N=(µ_pos_-µ_neg_)/((δ_pos_)x2+ (δ_neg_)x2)x0.5.

Briefly, HEK 293T cell were seeded in 15 cm dishes (5×10^6^ cells/dish) using DMEM supplemented with 10% FBS. After 24 hrs (day 2) of incubation (37ºC, 5% CO_2_), cells were transfected using PEI with either Jun-Nt, Fos-Ct, NiV F and G (3 µg of per plasmid) or mock transfected as described previously. On day 3, cells were counted, mixed into the positive and assay samples (Jun-Nt, Fos-Ct and NiV F and G expressing cells) and the negative controls (Jun-Nt, Fos-Ct and mock transfected cells) and seeded into 96 well plates (3.5×10^4^ cells/well in 100µl of media). In each mixture equal amounts of cells transfected with the different plasmids were included. Next, 2µL of a 50 µM compound stock (in 5% DMSO) was delivered into the cells (final concentration of 0.1% DMSO) using a MultiChannel Arm MCA 96 (Tecan). We used a Prestwick Chemical library (Sigma) containing 1120 compounds (note: currently the commercially available library contains 1280 compounds). In each plate 8 wells were treated just with DMSO (negative control) while 8 wells were reserved for the negative controls (see above). After 48 hours of incubation, fluorescence was measured directly on the plates in a EnSight multiplate reader (Perking Elmer). Results obtained from the screen were standardized using the Z-Score, calculated as follows: Z-Score=(x-μ)/δ, where x is the raw signal, μ is)/δ, where x is the raw signal, μ)/δ, where x is the raw signal, μ is is the mean signal and δ is the standard deviation of all the compound-containing wells of one plate indicates how many standard deviations a particular compound is above or below the mean of the plate. Primary hits were identified by calculating a Z-score for each compound and applying hit selection criteria (Z-Score <2).

To corroborate the anti-fusion activity of the selected compounds the experiment was repeated. Additionally, we measured the toxicity of the drugs in the exact same format by determining the number of viable cells utilizing the CellTiter 96 AQueous Non-Radioactive Cell Proliferation Assay (Promega) following the manufacturer instructions. Only those compounds with a Z-Score <2 in both rounds of the fusion assay and with a cell viability above 90 % (compared to the vehicle treated samples) were selected as hits and thus included in table 1.

**Table 1.** Inhibitors of NiV F and G induced cell-cell fusion.

| Name | Formula | MW | Previously reported as inhibitor of: |
| --- | --- | --- | --- |
| Anisomycin | C<sub>14</sub>H<sub>19</sub>NO<sub>4</sub> | 265.31 | Protein synthesis [37] |
| Cephaeline dihydrochloride heptahydrate | C<sub>28</sub>H<sub>54</sub>Cl<sub>2</sub>N<sub>2</sub>O<sub>11</sub> | 665.66 | Protein synthesis [38] |
| Digitoxigenin | C<sub>23</sub>H<sub>34</sub>O<sub>4</sub> | 374.53 | HSV [39] |
| Digoxin | C<sub>41</sub>H<sub>64</sub>O<sub>14</sub> | 780.96 | HSV [39] |
| Strophantine octahydrate | C<sub>29</sub>H<sub>60</sub>O<sub>20</sub> | 728.79 | HSV [39] |
| Emetine dihydrochloride | C<sub>29</sub>H<sub>42</sub>Cl<sub>2</sub>N<sub>2</sub>O<sub>4</sub> | 553.58 | Protein synthesis [40] |
| Proscillaridin A | C<sub>30</sub>H<sub>42</sub>O<sub>8</sub> | 530.66 | HBV [41] |

## 3. Results

### 3.1 Assay design

We sought to design a methodology for the study of cell-cell fusion triggered by viral proteins. For this porpoise, we exploited the bimolecular complementation properties of the Venus Fluorescent Protein (VFP) [22,26]. The reconstitution of a spitted VFP, a technique known as BiFC (Bimolecular Fluorescence Complementation) assay, has been extensively used for more than a decade for the identification and investigation of protein-protein interactions, even in genome wide analysis [27]. Briefly, the VFP can be divided into two non-fluorescent fragments (amino terminal (Nt) and carboxy terminal (Ct)). Reconstitution of the VFP native structure, and thus fluorescence, from these two fragments can be achieved if the Nt and Ct halves are brought into close proximity. This approximation might be prompted by the fusion of the VFP Nt and Ct fragments to two proteins or domains that spontaneously form a dimer, eg. Jun and Fos [28] (Figure 1A).

**Figure 1.**
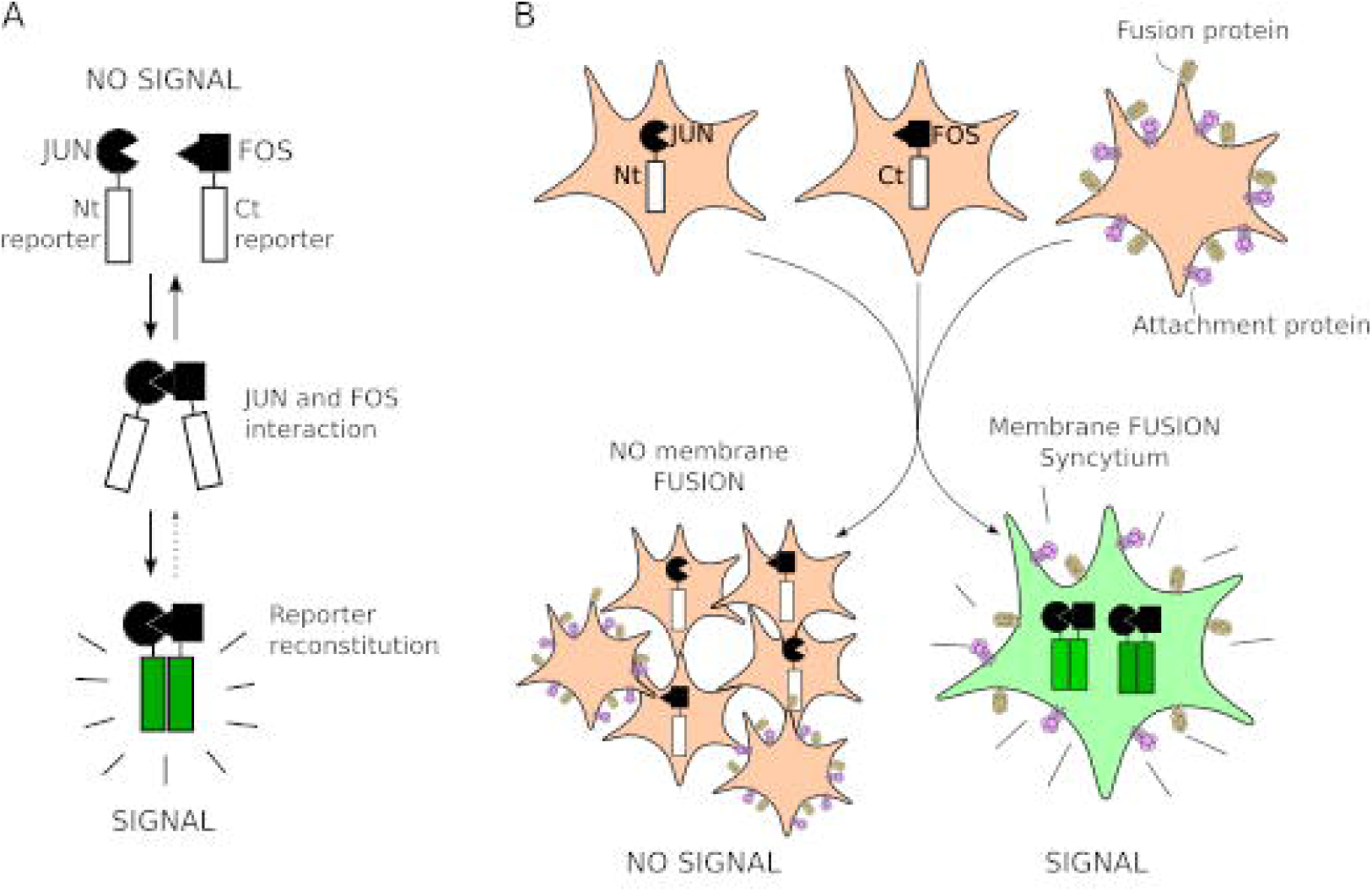
Schematic representation of BiMuC. **A**. Representation of the Bimolecular Complementation. **B**. Schematic representation of the Bimolecular Multicellular Complementation assay.

To analyze the cell-cell fusion, the Nt and Ct VFP segments fused to their corresponding interaction partners could be expressed in two independent cell pools while the viral proteins required for the formation of the syncytium are transfected into a third pool of cells (Figure 1B). After mixing these three cell populations, a successful fusion of the cell membranes by the viral proteins will bring the Nt and Ct chimeras together and the fluorescence of the VFP will be reconstituted. Moreover, if there is no syncytium no signal will be retrieve since the VFP fragments remain in different cells (Figure 1B). This methodology would tremendously facilitate the study of cell-cell fusion process in cells, which nowadays is primarily done by syncytium identification through microscopy observation. Furthermore, it could be applied for the identification of inhibitors of viral triggered membrane fusion and thus viral infection.

### 3.2 Assay validation

To test our hypothesis we fused the VFP Nt and Ct segments (as described in [22,29–31]) to Jun and Fos respectively [28], creating the Jun-Nt and Fos-Ct chimeras. These two constructs were independently transfected into HEK 293T human derived cells. Alongside, a third pool of cells was transfected with the NiV fusion (F) and attachment (G) proteins, which have been reported to be sufficient for cell-cell fusion [32,33] or mock transfected. A schematic representation of the designed protocol can be found in Figure 2. After transfection, cells were mixed into two groups both containing Jun-Nt and Fos-Ct expressing cells. Additionally, the positive group (+) was supplemented with cells expressing NiV F and G proteins while in the negative control (-) mock transfected cells were included. After 24 hrs, the fluorescence of both groups was measured. The results showed a staggering difference between samples with or without the viral proteins (Figure 3A). Only those samples that received the viral proteins (+), and thus formed a syncytium, were able to reconstitute the VFP. As expected, a fluorescence microscopy analysis revealed that cells that undergo cell-cell fusion are fluorescent while single cells remain non-fluorescents (Figure 3B). These results confirm the potential of our methodology for the analysis of cell-cell fusion events through the complementation of the VFP.

**Figure 2.**
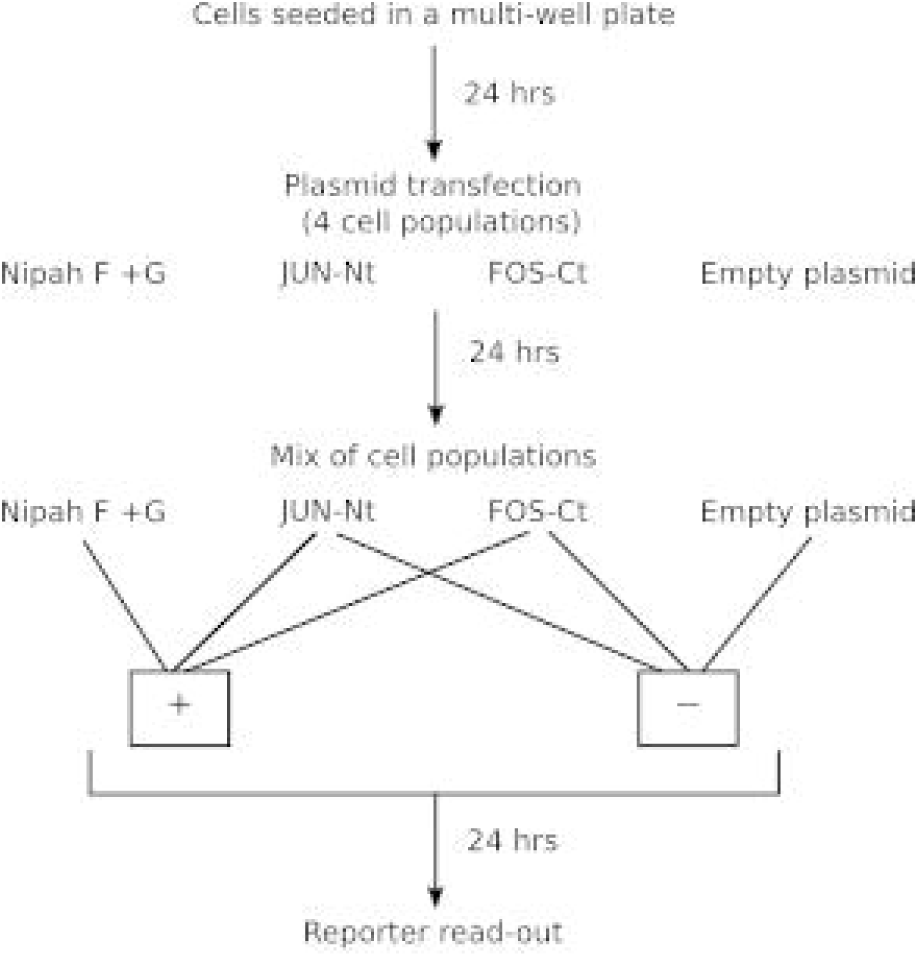
Schematic representation of the BiMuC assay workflow. The square with a + symbol in it denotes the positive control while the box with the – indicates the negative control. The cell pools included in both + and – controls are depicted.

**Figure 3.**
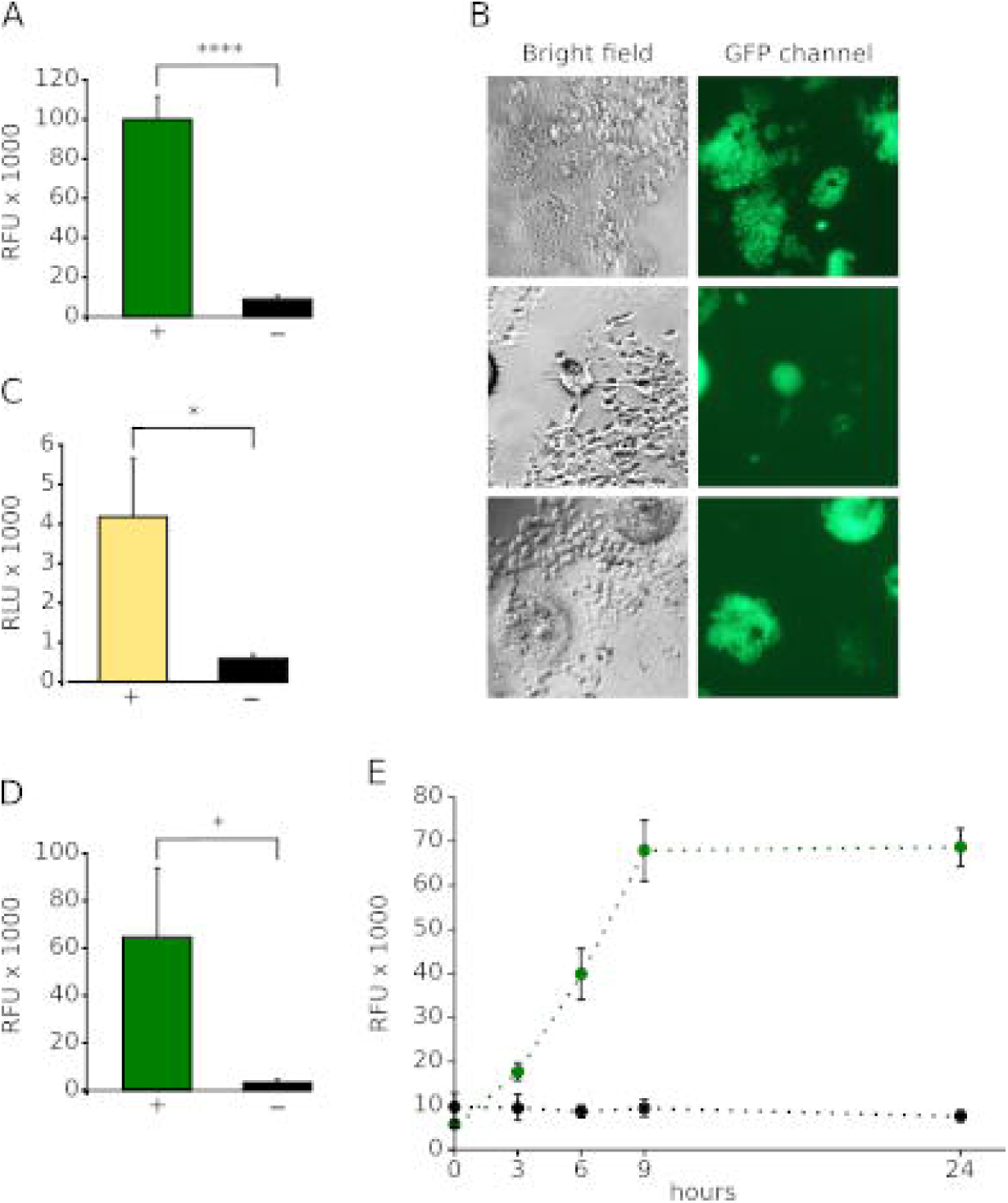
BiMuC assay validation. **A** Bar graph representing the fluorescence measurements results of positive (+, green including the NiV G and F, the Jun-Nt VFP and the Fos-Ct VFP cell pools) and negative (-, black, Jun-Nt VFP, Fos-Ct VFP, and mock transfected cells) controls. RFU, Relative Fluorescence Units. **B**. Bright and GFP channel fluorescence micrographies of positive samples. As expected, only fused cells can reconstitute the GFP signal. **C**. Bar graph showing the relative luminescence units (RLU) of the positive (+) and negative (-) for the BiMuC assay in which the luciferase has been use as a reporter. * p-value < 0.05, **** p-value< 0.001. **D**. results of the 2 pool approach using VFP as a reporter. Once again the positive control (+) is depicted with a green bar while the negative control (-) is shown using a black bar. **E**. Kinetics of the fusion process. RFU at multiple times (hours) post cell pool combination for the positive (green) and negative (black) controls. Error bars denote standard deviation of at least three replicates.

Many enzymes have shown the same reconstitution properties as the VFP [34]. For this reason, we decided to test whether the proposed methodology could be used with the luciferase (a widely used light producing enzyme) as a syncytium formation reporter. Similarly to the former approach, we fused Jun and Fos to the Nt and Ct ends of the firefly luciferase [35]. Next, we repeated the experiment as previously described (Figure 2) but using the new luciferase-based reporter. The results (Figure 3C) demonstrated that the luciferase is tolerated as a reporter by the system, which further increase our confidence in the devised methodology. We believe that the flexibility of the assay regarding the reporter increments the potential number of applications for our method.

Alternatively, we designed a protocol in which only 2 cell pools are required for the assay. In this case, one set of cells were transfected with Jun-Nt, NiV F and G proteins while the other received just Fos-Ct. Once the cells were mixed and the syncytium occurred a strong fluorescent signal was observed (Figure 3D). Contrarily, mock transfected cells remained non-fluorescent. This methodological variation represents a simplified but equally efficient version of the previously described assay. However, due to the increased variability observed when using this variation of the method we decided to employ for subsequent analysis the original design in which three pools of cells are mixed.

Next, we decided to explore the kinetics of the fusion process. For this experiment we proceeded using the three pool approach and the VFP as a reported. This time the samples were incubated, after mixing the three cell pools, 3, 6, and 9 hours and the results compared with the 24 hour-incubation formerly tested (Figure 3E). Two main conclusions can be obtained with this assay. First, the fusion process (at the present conditions) is over after nine hours. Second, the methodology allows a kinetic analysis of this type of fusion process since the signal increases steadily throughout the assay until the 9 hour mark.

### 3.3 Comparative analysis of the fusion and attachment proteins from the Henipahvirus genus

Once we have established a method for the study of syncytium formation we decided to compare the cell-cell fusion abilities of the F and G proteins from the closely related NiV and HeV. For this experiment we utilized the VFP-based assay previously described. Samples receiving cells expressing Jun-Nt, Fos-Ct and NiV F and G proteins were used as a positive control (+) while those that incorporate Jun-Nt, Fos-Ct but did not include cells transfected with any fusion and attachment protein serviced as negative control (-) (Figure 4). Our results indicated that HeV F and G proteins (HF/HG) provoke cell-cell fusion, as was previously reported in the literature [36]. However, the fluorescence levels of the HF/HG samples were, approximately, half than those observed when NiV F and G proteins were used to prompt the syncytium formation.

**Figure 4.**
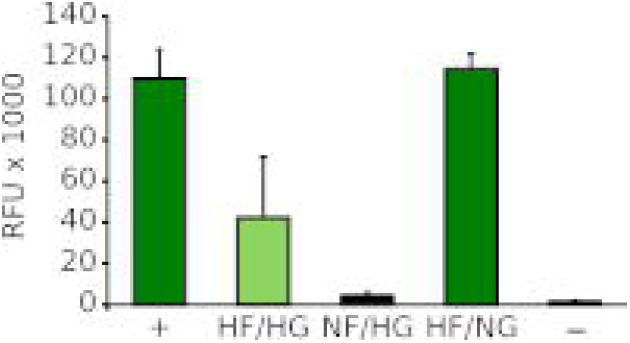
Comparative analysis of NiV and HeV fusion and attachment proteins. All samples were transfected with Jun-Nt VFP, Fos-Ct VFP except the negative control (-, mock transfected). Additionally, the positive control (+) was transfected with NiV F and G protein, HF/HG with HeV virus F and G proteins, NF/HG with NiV F and HeV G proteins, and HF/NG with HeV F and NiV G protein. Error bars denote standard deviation of at least three replicates.

Next, we decided to perform complementation studies between the fusion and attachment proteins of both viruses (NiV F and HeV G, NF/HG; and HeV F and NiV G, HF/NG). Surprisingly, the inclusion of cells expressing the NF/HG combination did not triggered fusion of cellular membranes while the HF/NG mix was successful stimulating the formation of a syncytium as it produced fluorescent levels comparable to the positive control. Suggesting that, despite of the similarity between the attachment proteins among both viruses, HeV G protein can not activate NiV F fusion activity.

### 3.4 A Bimolecular Multicelular complementation system for High-throughput small molecule identification

The sensitivity, simplicity and customization possibilities of the proposed methodology are ideal for the identification of new viral fusion and attachment proteins inhibitors. These small molecules screenings are performed in a high (or medium)-throughput compatible formats. Consequently, we decided to analyze the performance of our assay in 96 and 384 wells plate formats. The results (Figure 5), indicate that our assay is suitable for a 96 and even a 384 well format, regardless of whether we use 2 or 3 sets of cells.

**Figure 5.**
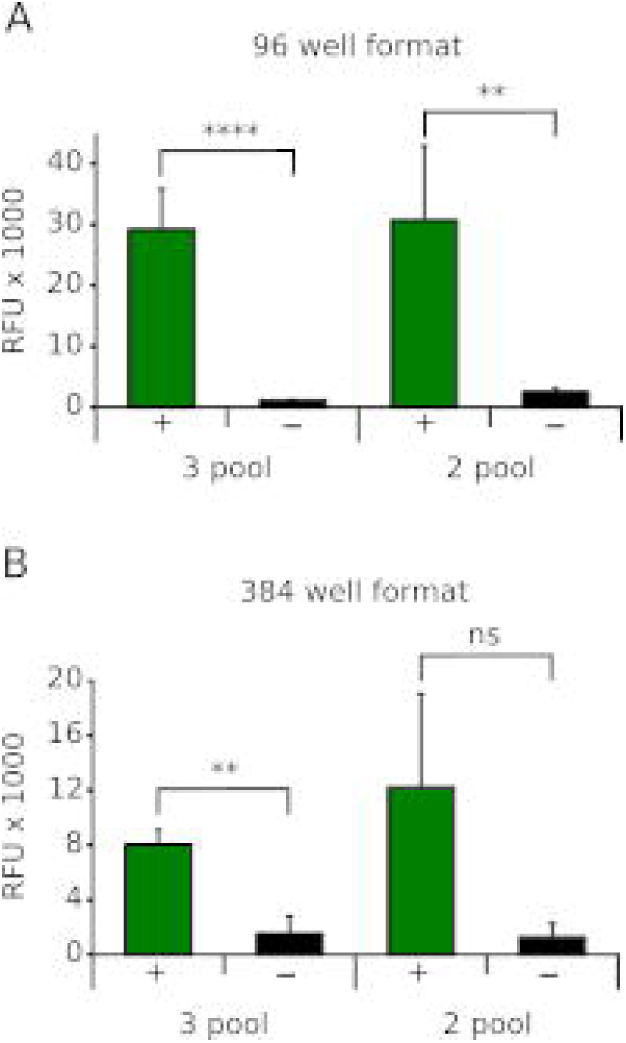
Miniaturization of the assay. **A**. BiMuC assay using the VFP as a reporter in a 96 (A) and 384 (B) well format. 3 pool and 2 pool indicates whether the 3 cell pools or the 2 cell pools approach was used. Positive control (+), negative control (-). Error bars denote standard deviation of at least three replicates.

Prompted by these results, we decided to perform a small molecules screening as a proof of concept. The assay was performed in a 96 well format at the *Screening laboratory of the Centro de Investigación Principe Felipe* (Valencia, Spain). We verified the suitability of our protocol for use in a High-throughput screening (HTS) by the Z’ factor (Z’=0.77) [25] and the signal-to-noise ratio (S/N=12.99). For the screening, 293T cells were seeded in complete media and 24 hrs later transfected with Jun-Nt, Fos-Ct, NiV F and G or mock transfected (Figure 6). After transfection, cells were mixed as described previously and treated with the appropriated small molecule. In each plate, 8 wells were reserved as a positive control (treated with DMSO) and 8 wells as a negative control (did not contain NiV F and G expressing cells). Next, cells were incubated 48 hrs allowing the small molecule to act, followed by fluorescence measurement and hit selection on an decreased signal over the vehicle-treated cells.

**Figure 6.**
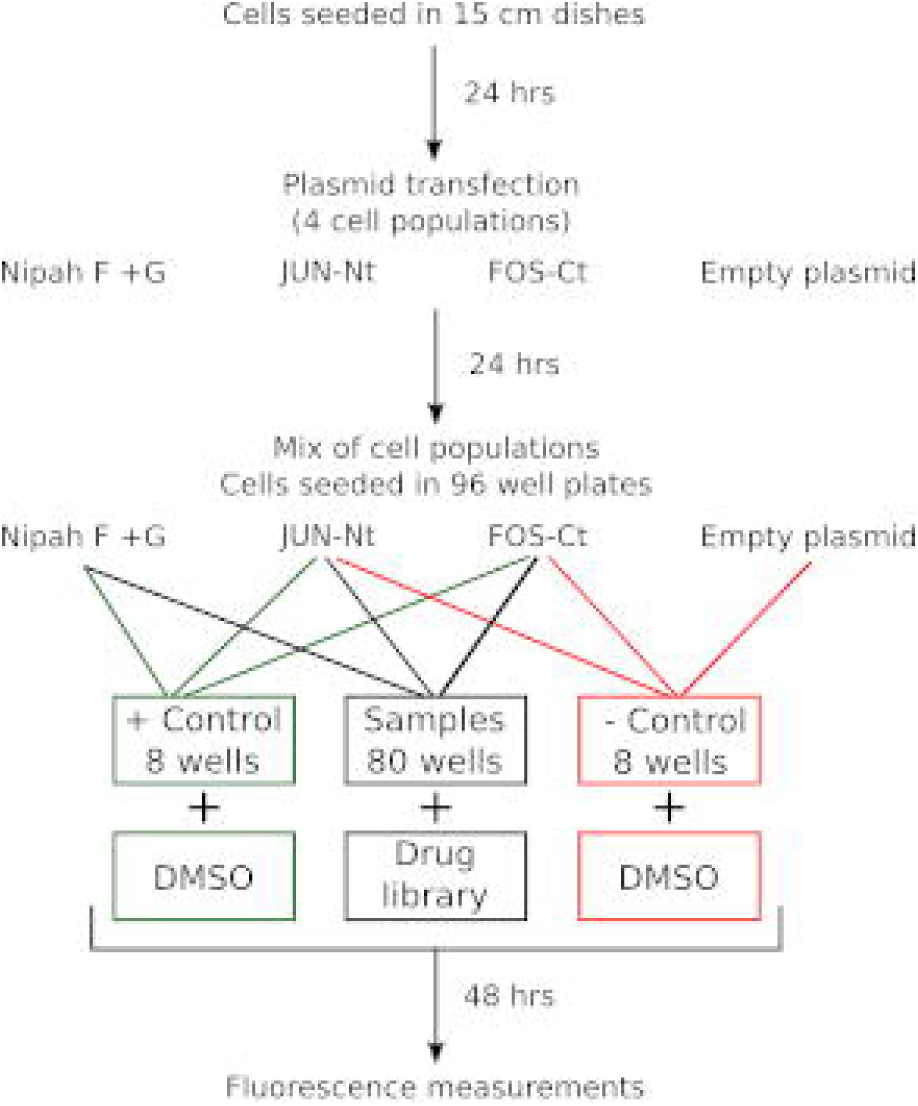
Schematic representation of the BiMuC assay high-throughput workflow. Representation of the workflow used to identify NiV fusion inhibitors.

A library consisting of 1120 chemically diverse small molecules, most of them approved drugs, (Prestwick Chemical) was screened and the results standardized by the Z-Score (primary hits were selected based on a Z-Score of <2). We identified 16 hits, which represent a hit rate of 0.014%. After a hit confirmation, using the same experimental conditions, and a cell proliferation assay where those compounds with a toxicity above 10% were eliminated the number of hits was reduced to 7 (hit rate = 0.0062%). Selected hits are included in table 1.

Many of the identified compounds were previously reported as entry inhibitors (Table 1) for Herpes simplex virus (HSV) [39] or Hepatitis B virus (HBV) [41]. Interestingly, these compounds are currently use as a cardiac stimulants [42]. Contrarily, those small molecules not identified as viral inhibitors are described in the literature as a protein synthesis inhibitors (see Table 1 for references).

## 4. Discussion

Viral infections have an enormous impact on global health. Just in the USA, it is estimated that influenza infections alone are responsible for 610,660 life lost and 3.1 million hospitalized days with an associated economic burden of $87.1 billion [43]. Thus, prevention and treatment of viral infection must be a priority. Treatment of viral infections generally often relies on anti-viral small molecules. Direct-acting antivirals (DTA), that is to say, antivirals that target the virus life cycle, are tremendously efficacious on blocking the viral replication [44]. Unfortunately, we have a very limited range of DTA at our disposal. Furthermore, some of the current blockbusters on viral inhibition might not remain so due to apparition of viral resistance [45]. Therefore, we must keep working on the development of new and improved antivirals. However, discovery of DTA has proven challenging at best, particularly, for BSL3 and 4 pathogens due to the restrictions imposed by the bio-safety measures. To facilitate the development of new entry inhibitors we have designed a new methodology that allows the quantification of cell-cell fusion events in a simple manner.

Our protocol, based on the bimolecular complementation properties of some protein reporters such as the VFP or the Renilla luciferase, facilitates the identification and quantification of cell-cell fusion events without the assistance of a microscope-base technology. Our results clearly indicate that this approach is robust and reproducible (p-value < 0.0001 was observed between the positive and negative samples on Figure 3A). Furthermore, our assay displayed a great flexibility. We could substituted the Venus fluorescent protein by the Renilla luciferase as a reporter or simplify the assay by using two cell pools instead of three and were still able to obtain solid results in both cases (Figure 3C and 3D). This flexibility is a great advantage, since every experiment has its own requirements the more adaptable the technique is the more application it might have. In theory, we could apply our approach not only to the study of viral induced cell-cell fusion (as we did for NiV and HeV viruses with great success) but also to any other cellular process in which a syncytium is formed, eg. the formation of the muscle fibers.

We also utilized the described methodology for the identification of small molecules capable of inhibiting the syncytium formation driven by the expression of NiV F and G proteins. In our screening we identified several steroidal glycosides, currently use for the treatment of cardiac disorders, that were previously described as HSV and HBV inhibitors [39]. This class of drugs, inhibitors of Na^+^/K^+^-ATPase, were thought to act at the early stage of HSV replication and the virus release stage but not to block virus entry or attachment [39]. Nonetheless, some other type of ATPase inhibitors, such as concanamycin A and SS33410, have been reported to inhibit poliovirus entry [46]. However, in this case, since poliovirus is a non enveloped virus, the reports refinfluenza to the endocytosis of the viral particle. Collectively, all the accumulate data regarding steroidal glycosides indicate that membrane rearrangements either the formation of endocytic vesicles of the fusion of viral and cellular membranes requires a intact concentration of ions across the cellular membrane but much work must be done to fully understand the process behind the mechanism of inhibition.

In our primary screen we identified 16 hits, which represent a hit rate of 0.014%. Smaller than the average hit rate for a HTS [47-50]. This is particularly striking since our assay was done in singles an not in duplicates like many other screenings. Nonetheless, the reduced hit rate observed is a welcome characteristic. Many HTS suffer from an elevate rate of false positives identification. Consequently, some steps must be introduced in the experimental protocol to reduce/eliminate the presence of the afore mentioned false positives. Therefore, the reduced hit rate of a robust assay like ours, not only simplifies the protocol, but also reduces the time and costs associated with the development of a new antiviral drug.

In conclusion, we have developed a new, simple, versatile, and safe methodology for the identification and quantification of syncytium formation applicable in small scale for an in-depths analysis of the cell-cell fusion process and in large scale to HTS small molecule identification of viral inhibitors.

## Acknowledgments and Funding

We would like to thank Dr. M. Shaw laboratory and Dr. BH Lee (Ichan School of Medicine at Mount Sinai) for the NiV and HeV F and G plasmids respectively. This work was supported by grants from the Spanish Ministry of Economy and Competitiveness (MINECO) (grant no. BFU2016-79487), and from the Generalitat Valenciana (GV/2016/139, Program Grupos Emergentes; and PROMETEOII/2014/061, Program Grupos de Excelencia)

## References

[1] Palese P, Shaw ML. Fields Virology. Orthomyxoviridae Viruses Their Replication 5th Edn Phila PA Lippincott Williams Wilkins Wolters Kluwer Bus 2007:1647–1689.

[2] Rey FA, Lok S-M. Common Features of Enveloped Viruses and Implications for Immunogen Design for Next-Generation Vaccines. Cell 2018;172:1319–34. doi:10.1016/j.cell.2018.02.054.

[3] Hidari KIPJ, Abe T, Suzuki T. Carbohydrate-related inhibitors of dengue virus entry. Viruses 2013;5:605–18. doi:10.3390/v5020605.

[4] Araújo LAL, Almeida SEM. HIV-1 diversity in the envelope glycoproteins: implications for viral entry inhibition. Viruses 2013;5:595–604. doi:10.3390/v5020595.

[5] Sun Z, Pan Y, Jiang S, Lu L. Respiratory syncytial virus entry inhibitors targeting the F protein. Viruses 2013;5:211–25. doi:10.3390/v5010211.

[6] Yang J, Li M, Shen X, Liu S. Influenza A virus entry inhibitors targeting the hemagglutinin. Viruses 2013;5:352–73. doi:10.3390/v5010352.

[7] De Feo CJ, Weiss CD. Escape from human immunodeficiency virus type 1 (HIV-1) entry inhibitors. Viruses 2012;4:3859–911.

[8] Yu F, Lu L, Du L, Zhu X, Debnath AK, Jiang S. Approaches for identification of HIV-1 entry inhibitors targeting gp41 pocket. Viruses 2013;5:127–49. doi:10.3390/v5010127.

[9] Markowitz M, Meyers K. Extending access with long-acting antiretroviral therapy: the next advance in HIV-1 therapeutics and prevention. Curr Opin HIV AIDS 2015;10:216–8. doi:10.1097/ COH.0000000000000167.

[10] Kesiraju S, Srikanti P, Sahariah S. Hepatitis C infection in renal transplantation: pathogenesis, current impact and emerging trends. Virusdisease 2017;28:233–41. doi:10.1007/s13337-017-0393-5.

[11] Rumlová M, Ruml T. In vitro methods for testing antiviral drugs. Biotechnol Adv 2017. doi:10.1016/j.biotechadv.2017.12.016.

[12] Pasieka TJ, Woolson RF, Grose C. Viral induced fusion and syncytium formation: measurement by the Kolmogorov–Smirnov statistical test. J Virol Methods 2003;111:157–61. doi:10.1016/S0166-0934(03)00152-6.

[13] Kramer S, Buontempo P, Agrawal S, Ralston R. Imaging-based assay for identification and characterization of inhibitors of CXCR4-tropic HIV-1 envelope-dependent cell-cell fusion. J Biomol Screen 2011;16:668–75. doi:10.1177/1087057111403480.

[14] Pine PS, Weaver JL, Oravecz T, Pall M, Ussery M, Aszalos A. A semiautomated fluorescence-based cell-to-cell fusion assay for gp120-gp41 and CD4 expressing cells. Exp Cell Res 1998;240:49–57. doi:10.1006/excr.1998.3939.

[15] Hong YL, Wu LH, Cui M, McMaster G, Hunt SW, Chung FZ. New reporter cell lines to study macrophage-tropic HIV envelope protein-mediated cell-cell fusion. AIDS Res Hum Retroviruses 1999;15:1667–72. doi:10.1089/088922299309702.

[16] Chiba H, Asanuma S, Okamoto M, Inokoshi J, Tanaka H, Fujita K, et al. A simple screening system for anti-HIV drugs: syncytium formation assay using T-cell line tropic and macrophage tropic HIV env expressing cell lines--establishment and validation. J Antibiot (Tokyo) 2001;54:818–26.

[17] Lifson JD, Feinberg MB, Reyes GR, Rabin L, Banapour B, Chakrabarti S, et al. Induction of CD4-dependent cell fusion by the HTLV-III/LAV envelope glycoprotein. Nature 1986;323:725–8. doi:10.1038/323725a0.

[18] Cheng D-C, Zhong G-C, Su J-X, Liu Y-H, Li Y, Wang J-Y, et al. A sensitive HIV-1 envelope induced fusion assay identifies fusion enhancement of thrombin. Biochem Biophys Res Commun 2010;391:1780–4. doi:10.1016/j.bbrc.2009.12.155.

[19] Edinger AL, Doms RW. A Cell-Cell Fusion Assay to Monitor HIV-1 Env Interactions with Chemokine Receptors. Methods Mol Med 1999;17:41–9. doi:10.1385/0-89603-369-4:41.

[20] Herschhorn A, Finzi A, Jones DM, Courter JR, Sugawara A, Iii ABS, et al. An Inducible Cell-Cell Fusion System with Integrated Ability to Measure the Efficiency and Specificity of HIV-1 Entry Inhibitors. PLOS ONE 2011;6:e26731. doi:10.1371/journal.pone.0026731.

[21] Bradley J, Gill J, Bertelli F, Letafat S, Corbau R, Hayter P, et al. Development and automation of a 384-well cell fusion assay to identify inhibitors of CCR5/CD4-mediated HIV virus entry. J Biomol Screen 2004;9:516–24. doi:10.1177/1087057104264577.

[22] Kerppola TK. Design and implementation of bimolecular fluorescence complementation (BiFC) assays for the visualization of protein interactions in living cells. Nat Protoc 2006;1:1278–86. doi:10.1038/nprot.2006.201.

[23] Paulmurugan R, Gambhir SS. Monitoring protein-protein interactions using split synthetic renilla luciferase protein-fragment-assisted complementation. Anal Chem 2003;75:1584–9.

[24] Islam MA, Park T-E, Singh B, Maharjan S, Firdous J, Cho M-H, et al. Major degradable polycations as carriers for DNA and siRNA. J Control Release Off J Control Release Soc 2014;193:74–89. doi:10.1016/j.jconrel.2014.05.055.

[25] Zhang J, Chung T, Oldenburg K. A Simple Statistical Parameter for Use in Evaluation and Validation of High Throughput Screening Assays. J Biomol Screen 1999;4:67–73. doi:10.1177/108705719900400206.

[26] Magliery TJ, Wilson CGM, Pan W, Mishler D, Ghosh I, Hamilton AD, et al. Detecting protein-protein interactions with a green fluorescent protein fragment reassembly trap: scope and mechanism. J Am Chem Soc 2005;127:146–57. doi:10.1021/ja046699g.

[27] Miller KE, Kim Y, Huh W-K, Park H-O. Bimolecular Fluorescence Complementation (BiFC) Analysis: Advances and Recent Applications for Genome-Wide Interaction Studies. J Mol Biol 2015;427:2039–55. doi:10.1016/j.jmb.2015.03.005.

[28] van Dam H, Castellazzi M. Distinct roles of Jun: Fos and Jun: ATF dimers in oncogenesis. Oncogene 2001;20:2453–64. doi:10.1038/sj.onc.1204239.

[29] Kerppola TK. Bimolecular fluorescence complementation (BiFC) analysis as a probe of protein interactions in living cells. Annu Rev Biophys 2008;37:465–87. doi:10.1146/annurev.biophys.37.032807.125842.

[30] Hu C-D, Chinenov Y, Kerppola TK. Visualization of interactions among bZIP and Rel family proteins in living cells using bimolecular fluorescence complementation. Mol Cell 2002;9:789–98.

[31] Ghosh I, Hamilton AD, Regan L. Antiparallel Leucine Zipper-Directed Protein Reassembly: Application to the Green Fluorescent Protein. J Am Chem Soc 2000;122:5658–9. doi:10.1021/ja994421w.

[32] Vera-Velasco NM, García-Murria MJ, Sánchez Del Pino MM, Mingarro I, Martinez-Gil L. Proteomic composition of Nipah virus-like particles. J Proteomics 2018;172:190–200. doi:10.1016/ j.jprot.2017.10.012.

[33] Chua KB, Bellini WJ, Rota PA, Harcourt BH, Tamin A, Lam SK, et al. Nipah virus: a recently emergent deadly paramyxovirus. Science 2000;288:1432–5.

[34] Wehr MC, Rossner MJ. Split protein biosensor assays in molecular pharmacological studies. Drug Discov Today 2016;21:415–29. doi:10.1016/j.drudis.2015.11.004.

[35] Fujikawa Y, Kato N. TECHNICAL ADVANCE: Split luciferase complementation assay to study protein–protein interactions in Arabidopsis protoplasts. Plant J 2007;52:185–95. doi:10.1111/j.1365-313X.2007.03214.x.

[36] Hooper P, Zaki S, Daniels P, Middleton D. Comparative pathology of the diseases caused by Hendra and Nipah viruses. Microbes Infect 2001;3:315–22.

[37] Pubchem. Anisomycin n.d. https://pubchem.ncbi.nlm.nih.gov/compound/253602 (accessed April 24, 2018).

[38] Pubchem. Cephaeline n.d. https://pubchem.ncbi.nlm.nih.gov/compound/442195 (accessed April 24, 2018).

[39] Su C-T, Hsu JT-A, Hsieh H-P, Lin P-H, Chen T-C, Kao C-L, et al. Anti-HSV activity of digitoxin and its possible mechanisms. Antiviral Res 2008;79:62–70. doi:10.1016/j.antiviral.2008.01.156.

[40] Pubchem. Emetine hydrochloride n.d. https://pubchem.ncbi.nlm.nih.gov/compound/3068143 (accessed April 24, 2018).

[41] Okuyama-Dobashi K, Kasai H, Tanaka T, Yamashita A, Yasumoto J, Chen W, et al. Hepatitis B virus efficiently infects non-adherent hepatoma cells via human sodium taurocholate cotransporting polypeptide. Sci Rep 2015;5:17047. doi:10.1038/srep17047.

[42] Belz GG, Breithaupt-Grögler K, Osowski U. Treatment of congestive heart failure--current status of use of digitoxin. Eur J Clin Invest 2001;31 Suppl 2:10–7.

[43] Molinari N-AM, Ortega-Sanchez IR, Messonnier ML, Thompson WW, Wortley PM, Weintraub E, et al. The annual impact of seasonal influenza in the US: measuring disease burden and costs. Vaccine 2007;25:5086–96. doi:10.1016/j.vaccine.2007.03.046.

[44] Aghemo A, Piroth L, Bhagani S. What do clinicians need to watch for with direct-acting antiviral therapy? J Int AIDS Soc 2018;21 Suppl 2:e25076. doi:10.1002/jia2.25076.

[45] Zuo X, Huo Z, Kang D, Wu G, Zhou Z, Liu X, et al. Current insights into anti-HIV drug discovery and development: a review of recent patent literature (2014-2017). Expert Opin Ther Pat 2018;28:299–316. doi:10.1080/13543776.2018.1438410.

[46] Irurzun A, Carrasco L. Entry of poliovirus into cells is blocked by valinomycin and concanamycin A. Biochemistry (Mosc) 2001;40:3589–600.

[47] Ortigoza MB, Dibben O, Maamary J, Martinez-Gil L, Leyva-Grado VH, Abreu P, et al. A novel small molecule inhibitor of influenza A viruses that targets polymerase function and indirectly induces interferon. PLoS Pathog 2012;8:e1002668. doi:10.1371/journal.ppat.1002668.

[48] Meleza C, Thomasson B, Ramachandran C, O’Neill JW, Michelsen K, Lo M-C. Development of a scintillation proximity binding assay for high-throughput screening of hematopoietic prostaglandin D2 synthase. Anal Biochem 2016;511:17–23. doi:10.1016/j.ab.2016.07.028.

[49] Weisshaar M, Cox R, Morehouse Z, Kumar Kyasa S, Yan D, Oberacker P, et al. Identification and Characterization of Influenza Virus Entry Inhibitors through Dual Myxovirus High-Throughput Screening. J Virol 2016;90:7368–87. doi:10.1128/JVI.00898-16.

[50] Martínez-Gil L, Ayllon J, Ortigoza MB, García-Sastre A, Shaw ML, Palese P. Identification of small molecules with type I interferon inducing properties by high-throughput screening. PloS One 2012;7:e49049. doi:10.1371/journal.pone.0049049.

